# DPHL v2: An updated and comprehensive DIA pan-human assay library for quantifying more than 14,000 proteins

**DOI:** 10.1101/2023.01.07.523067

**Authors:** Zhangzhi Xue, Tiansheng Zhu, Fangfei Zhang, Cheng Zhang, Nan Xiang, Liujia Qian, Xiao Yi, Yaoting Sun, Wei Liu, Xue Cai, Linyan Wang, Xizhe Dai, Liang Yue, Lu Li, Thang V. Pham, Sander R. Piersma, Qi Xiao, Meng Luo, Cong Lu, Jiang Zhu, Yongfu Zhao, Guangzhi Wang, Junhong Xiao, Tong Liu, Zhiyu Liu, Yi He, Qijun Wu, Tingting Gong, Jianqin Zhu, Zhiguo Zheng, Juan Ye, Yan Li, Connie R. Jimenez, A Jun, Tiannan Guo

## Abstract

A comprehensive pan-human spectral library is critical for biomarker discovery using mass spectrometry (MS)-based proteomics. DPHL v1, a previous pan-human library built from 1096 data-dependent acquisition (DDA) MS data of 16 human tissue types, allows quantifying 10,943 proteins. However, a major limitation of DPHL v1 is the lack of semi-tryptic peptides and protein isoforms, which are abundant in clinical specimens. Here, we generated DPHL v2 from 1608 DDA-MS data acquired using Orbitrap mass spectrometers. The data included 586 DDA-MS newly acquired from 17 tissue types, while 1022 files were derived from DPHL v1. DPHL v2 thus comprises data from 24 sample types, including several cancer types (lung, breast, kidney, and prostate cancer, among others). We generated four variants of DPHL v2 to include semi-tryptic peptides and protein isoforms. DPHL v2 was then applied to a publicly available colorectal cancer dataset with 286 DIA-MS files. The numbers of identified and significantly dysregulated proteins increased by at least 21.7% and 14.2%, respectively, compared with DPHL v1. Our findings show that the increased human proteome coverage of DPHL v2 provides larger pools of potential protein biomarkers.

## Introduction

Mass spectrometry (MS)-based quantitative proteomics is widely used for protein biomarker discovery^1–3^. The subsequent biomarker validation is often performed with targeted proteomics methods, such as selected reaction monitoring (SRM)^4^ and parallel reaction monitoring (PRM)^5^. Recently, biomarker discovery and validation have been increasingly performed with targeted analysis of data-independent acquisition (DIA) MS data^6^, an emerging strategy for high-throughput proteomics analyses with a high level of reproducibility^7^. A spectral library containing experimental peptide precursor information is crucial for SRM- and PRM-based protein biomarker validation, as well as DIA-based biomarker discovery^7^. In recent years, spectral libraries have been established for several organisms, such as human^8, 9^, mouse^10^, zebrafish^11^, *Arabidopsis thaliana*^12^, and *Escherichia coli*^13^. To support the identification of new protein biomarkers, the comprehensiveness of a spectral library is crucial.

The Human Proteome Project (HPP)^14^ launched by Human Proteome Organization (HUPO) has reported the community-based ten-year achievement of a high-stringency proteome blueprint of 17,874 Protein Evidence 1 (PE1) proteins in 2020, covering 90.4% of the human proteome^15^. A pan-human spectral library (PHL), containing 149,130 peptide precursors and 10,322 proteins, was developed to analyze Sequential Window Acquisition of All Theoretical Mass Spectra (SWATH-MS) data acquired on SCIEX TripleTOF Systems^8^. Another DIA pan-human library (DPHL v1) for Orbitrap data comprises 289,237 peptide precursors and 10,943 proteins^9^.

However, the proteins in these two libraries are proteotypic; protein isoforms are not included. The isoforms of each protein family may result from post-translational modifications, splice variants, proteolytic products, genetic variations, or somatic recombination occurring during protein evolution^16^, and participate in different biological processes^17^. Therefore, a specific protein isoform could be a valuable biomarker. A spectral library with significant coverage of the human proteome and its protein isoforms is thus needed. Additionally, previous studies demonstrated that only ~10-15% of all the tryptic peptides from a protein sample can be identified when about 50% of the protein identifications are based on a single tryptic peptide due to the intrinsic chemical properties of tryptic peptides^18–20^. Therefore, identifying more peptides (e.g., non-tryptic peptides), preferably at low computational costs, would increase the confidence in the proteins identified via tryptic peptides and increase the overall number of identifications.

Here, we present a large DIA spectral library (DPHL v2), generated from 24 different sample types and available in four variants. DPHL v2 includes more peptide precursors, peptides, and proteins than DPHL v1. It also provides higher coverage ratios, particularly for brain-, esophagus-, and ovary-specific or -enriched proteins, as well as FDA-approved drug targets. Two variants of DPHL v2 generated better identifications of the hallmark gene sets than DPHL v1. Finally, using a publicly available colorectal cancer (CRC) cohort, DPHL v2 provided larger numbers of protein and differentially expressed protein identifications than DPHL v1 and library-free method.

## Results and Discussion

### Data sources for generating DPHL v2

A total of 1608 raw MS data files were collected to build our spectral library. Among these, 586 files were newly generated from various samples, including tissue biopsies of prostate cancer (PCa), hepatocellular carcinoma (HCC), triple-negative breast cancer (TNBC), lung adenocarcinoma (LUAD), esophageal carcinoma, thyroid diseases, eyelid tumors, glioblastoma multiforme (GBM), healthy brain tissues, oral squamous cell carcinoma (OSCC), thymic diseases, ovarian cancer (OV), and cervix cancer. Additionally, blood plasma samples from acute myelocytic leukemia (AML), blood diseases, T-lineage acute lymphoblastic leukemia (T-ALL), and normal plasma exosome were included. Human chronic myelogenous leukemia cell line K562 was also included. Finally, the remaining 1022 files were derived from the DPHL v1 study by Zhu *et al*.^9^ The sample types and number of patients contributing to DPHL v2 are summarized in Figure 1A and Table S1.

**Figure 1.**
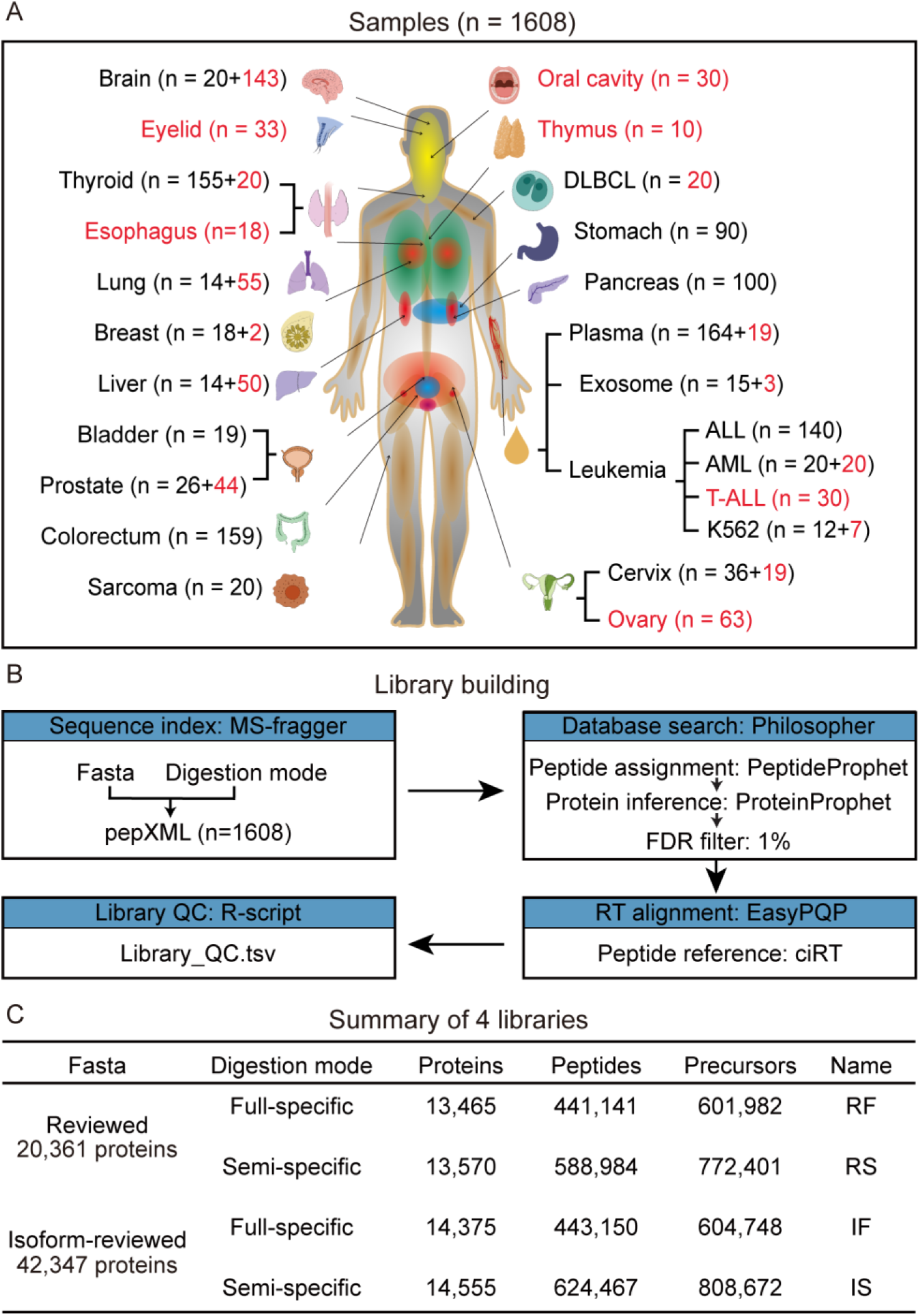
Sample types and workflow for building DPHL v2. (A) Number and type of samples included in this study. The ones that were missing from DPHL v1 are highlighted in red. (B) Computational pipeline for building DPHL v2. (C) Overview of the number of identified proteins, peptides, and precursors using our four library variants.

### Four variants of the pan-human spectral libraries

All the 1608 raw files were centroided and converted into mzXML as previously described^9^. These files were then combined to build our new spectral library. Two different annotation files (i.e., reviewed and isoform-reviewed fasta files) were used to search the mzXML spectra against two digestion modes (i.e., full-specific and semi-specific) using MS-Fragger (version 3.0)^21^. The reviewed fasta file was obtained from the UniProt database^22^ (accessed on 17 Jul. 2020); it included 20,361 reviewed human proteins and was used as the reference. The isoform-reviewed annotation file was also downloaded from UniProt (accessed on 5 Aug. 2020) and comprised 42,347 proteins, including 22,201 human isoforms. Philosopher^23^ (version 3.2.9) was used for library searching based on the spectra matches with a maximum of two missed cleavages and a false discovery rate < 0.01 for spectra, peptides, and proteins. By differently combining the two annotation files and the two digestion modes, we generated four library variants: RF (reviewed fasta sequence & full-specific digestion mode), RS (reviewed fasta sequence & semi-specific digestion mode), IF (isoform fasta sequence & full-specific digestion mode), IS (isoform fasta sequence & semi-specific digestion mode).

Next, in order to ensure the consistency of the results of different time gradients of the mass spectrum, we used EasyPQP (version 0.1.9, https://github.com/grosenberger/easypqp) to anchor the CiRT^21^ peptides for retention time (RT) normalization. Quality controls (QC) were then performed using an R script with the criteria next described to remove data of low quality. First, only precursors with multiple fragments (≥ 2) and a normalized RT range from −60 to 200 were retained. Second, fragments with a library intensity < 10 or a precursor charge of +1 were removed. Finally, peptides with only one precursor were retained. However, when a peptide had two precursors, we kept the one with the highest intensity if the absolute difference of the normalized RT between the two precursors was > 5; otherwise, both precursors were kept. When a peptide has more than two precursors, the averaged normalized RTs of all precursors and their differences with respect to their mean RT were calculated. Next, peptides with an absolute difference > 5 were excluded. When all the absolute values were > 5, the median normalized RT of all the precursors and their difference from the median RT were further calculated: only the peptides with a difference < 5 were then selected. The normalized RT correlations (+2/+3 states of each peptide) after these filtering steps are shown in Figure S1. Default parameters were used for all software unless otherwise indicated. The computational pipeline is schematized in Figure 1B.

### Characteristics of DPHL v2

We next evaluated DPHL v2 using DIALib-QC^24^ and found that all four variants of our pan-human spectral library are of high quality (Figure S2-5). We also characterized the four libraries in terms of peptide and protein identifications. As shown in Figure 1C, the RF library includes 601,982 peptide precursors, 441,141 peptides, and 13,465 proteins; the IF library includes 604,748 peptide precursors, 443,150 peptides, and 14,375 proteins. IS, another isoform-based library, comprises 808,672 peptide precursors, 624,467 peptides, and 14,555 proteins. Finally, the RS library contains 772,401 peptide precursors, 588,984 peptides, and 13,570 proteins. We then evaluated the protein identifications of the four libraries for each of the 24 sample types. As shown in Figures S6-7, the brain had the highest number of total and unique proteins among all sample types, possibly due to the larger number of brain tissues included (n = 163).

Next, we compared our four libraries with the PHL and DPHL v1 and found that our four libraries exhibited at least a 23.0% and 30.4% increase in protein coverage compared to DPHL v1 and the PHL, respectively. Among our four libraries, the isoform-based ones (IS and IF) comprise relatively high numbers of proteins (Figure 2A). Similarly, our four libraries exhibit considerably larger numbers of peptide (Figure 2C) and precursor (Figure 2E) identifications when compared to DPHL v1 and the PHL. In particular, the semi-specific digestion libraries (IS and RS) have the most significant numbers of peptide and precursor identifications. As shown in Figure 2B, 2D, and 2F, 7262 proteins, 89,328 peptides, and 103,704 precursors are shared among these six libraries, while 1,144 proteins, 165,041 peptides, and 253,673 precursors are shared only by our four libraries. These findings indicate that DPHL v2 provides higher coverage among precursors, peptides, and proteins than DPHL v1 and the PHL.

**Figure 2.**
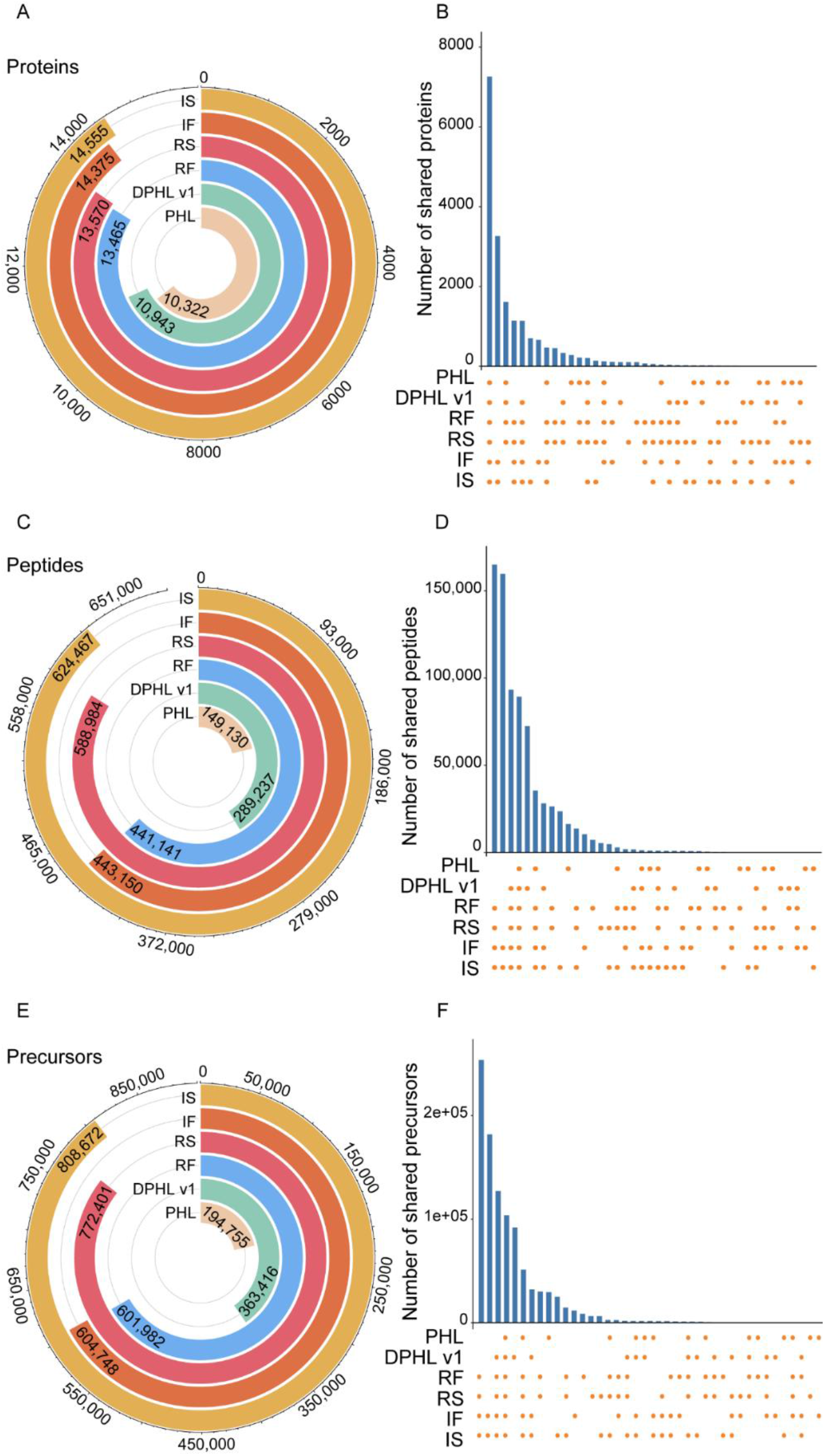
Comparison of the four variants of DPHL v2 (i.e., RF, RS, IF, and IS) with DPHL v1 and PHL. The circular bars show the protein (A), peptide (C), and precursor identifications (E) of the six libraries. The UpSet plots show the shared and unique protein (B), peptide (D), and precursor identifications (F) of the six libraries. PHL, pan-human spectral library; DPHL v1, DIA pan-human library generated by Zhu *et al*; RF, reviewed fasta sequence & full-specific digestion mode; RS, reviewed fasta sequence & semi-specific digestion mode; IF, isoform fasta sequence & full-specific digestion mode; IS, isoform fasta sequence & semi-specific digestion mode.

We next compared the numbers of shared proteins and peptides between our four library variants (*i.e*., between fasta files and digestion models) (Figure 3A, 3B). We found that protein identifications were affected mainly by the fasta file, while peptide identifications were affected by the digestion model. We also compared our four libraries with DPHL v1 in terms of the enriched/specific proteins from three tissues (brain, ovary, and esophagus; Figure 3C) obtained from the Human Protein Atlas (https://www.proteinatlas.org/, data available from v21.0.proteinatlas.org). Our results indicated that the coverages of our four libraries are superior to that of DPHL v1. Similarly, our four libraries provided higher coverage of FDA-approved drug targets than DPHL v1 (Figure 3C). In addition, the hallmark gene sets from the MSigDB v7.4 database (http://www.broad.mit.edu/gsea/msigdb/, accessed on 22 Nov. 2021)^25, 26^ were analyzed using these five libraries. We found that RF and RS cover more than 44% of the genes with well-defined biological states or processes, and both provide better coverages than DPHL v1 (Figure 3C). However, fewer coverages were found in the isoform-based libraries. One possible reason is that most genes from the hallmark gene sets are reviewed.

**Figure 3.**
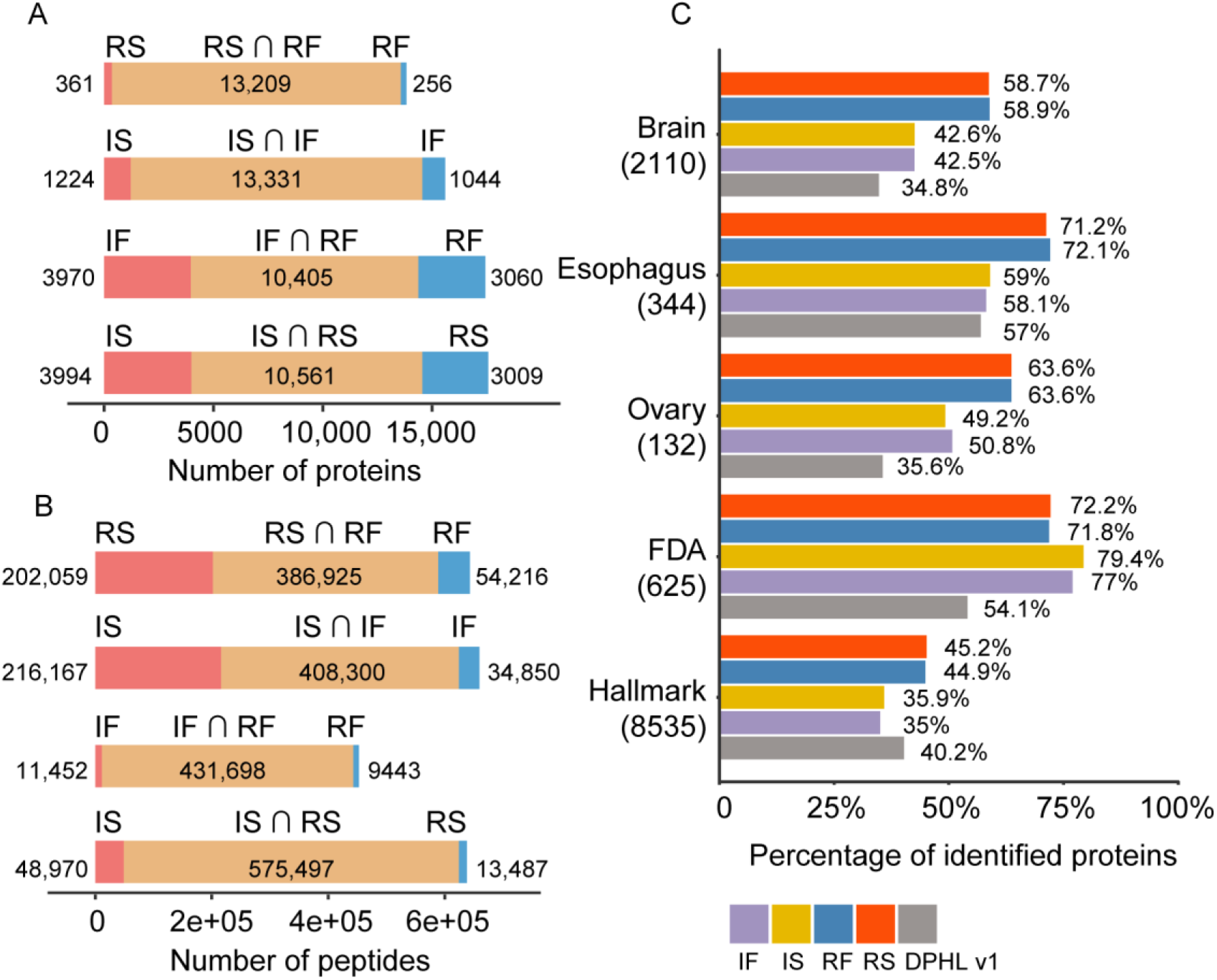
Comparison of the number of proteins (A) and peptides (B) identified with the same fasta sequence and the same digestion mode. (C) Percentage of proteins identified among DPHL v1 and our four libraries using hallmark gene sets, FDA-approved drug targets, and tissue-specific or tissue-enriched/enhanced proteins from brain, esophagus, and ovary samples.

### Applicability of DPHL v2 for DIA targeted data analysis

To assess the applicability of DPHL v2, we used our four libraries, DPHL v1, or a library-free method to analyze a CRC cohort, including 201 CRC cases, 40 benign samples, and 45 biological/technical replicates^27^. The missing values generated by our four libraries or DPHL v1 were comparable. On the other hand, the library-free method generated fewer missing values (Figure 4A). As shown in Figure 4B, the number of proteins identified with any variant of DPHL v2 was significantly higher than with DPHL v1 or the library-free method. A total of 978 proteins were identified by all six methods, while 166 were shared by our four libraries only (Figure 4C).

**Figure 4.**
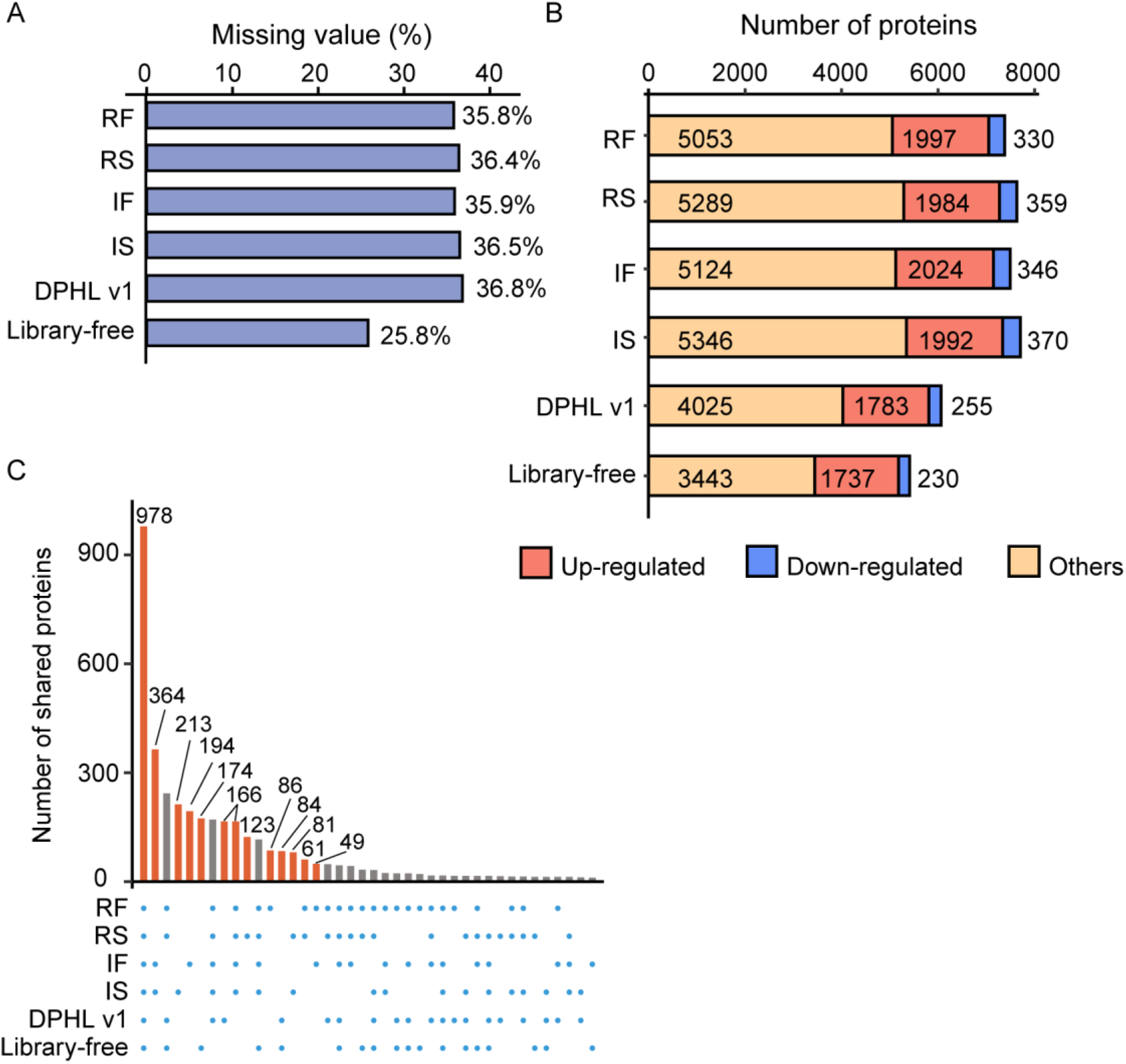
DIA analysis of CRC and benign samples. (A) Number of missing values obtained using the five libraries and library-free method. (B) Number of differentially expressed proteins between CRC and benign samples obtained using the five libraries and library-free method. Proteins with adjusted p-value < 0.01 and |FC| > 4 were selected as significantly differentially expressed. FC, fold change. (C) Protein identification overlaps across the six libraries.

In order to demonstrate the applicability of the library, we performed differential expression analyses of the CRC data generated using the six methods described above. Differential expressions were considered significant if their adjusted p-values were < 0.01 and their log_2_ (fold-change) absolute values were > 1. We obtained 1997 (RF), 1984 (RS), 2024 (IF), 1992 (IS), 1783 (DPHL v1), and 1737 (library-free) up-regulated (adjusted p-value < 0.01 & log_2_ (fold-change) > 1) proteins, and 330 (RF), 359 (RS), 346 (IF), 370 (IS), 255 (DPHL v1), and 230 (library-free) down-regulated (adjust p-value < 0.01 & log_2_ (fold-change) < −1) proteins (Figure 4B). Compared with the DPHLv1, the numbers of identified and significantly dysregulated proteins increased by at least 21.7% (RF) and 14.2% (RF). Compared with the analysis using only SwissProt reviewed proteins sequences, 463 and 472 differentially expressed protein isoforms were identified using IF and IS, respectively. Similarly, 94 and 92 proteins were dysregulated in the CRC tissues compared with the benign samples by semi-specific digestion modes. These findings show that DPHL v2 allows identifying a larger number of differentially expressed proteins or protein isoforms between tumors and benign samples, providing more options for subsequent investigations.

We next used our four libraries and DPHL v1 to analyze the CRC cohort using the sub-library strategy^27^, which refines a pan-human spectral library based on the tissue specificity. Compared with the conventional library search method, the sub-library strategy improved our results in all aspects (Figure S8A-C). First, the missing values were reduced by about 1% on average. The protein identifications increased by 22 (RF), 344 (RS), 103 (IF), 405 (IS), or 193 (DPHL v1). In the subsequent differential expression analysis, the total number of dysregulated proteins increased by 70 (RF), 163 (RS), 42 (IF), 203 (IS), and 20 (DPHL v1).

Finally, we built a random forest model based on the overlap dysregulated proteins generated by the four libraries to find new biomarkers. The 241 samples with 1426 proteins were randomly divided into the training set (N = 200) and the test set (N = 41). After a 5-fold cross validation, we identified 14 features that provided the highest accuracy for colorectal cancer, including S100A11, CEACAM6, GARS1, CDYL2, POTEKP, SCGN, SNCG, S100B, SCG2, NCAM1, OGN, CD81, COL28A1, CNRIP1 (Figure 5A). The area under the curve (AUC) of the training set and the test set achieved 1, 0.903 (Figure 5B), and the accuracy (ACC) achieved 0.988, 0.927, respectively (Figure 5C). Among these, S100A11^28, 29^, CEACAM6^30, 31^, CDYL2^32^, SCGN^33^, SNCG^34, 35^, S100B^36^, SCG2^37, 38^, NCAM1^39^, OGN^40^, CD81^41^, CNRIP1^42^, have been reported to be closely related to colorectal cancer. Three features (GARS1, POTEKP, COL28A1) may be new biomarkers for colorectal cancer.

**Figure 5.**
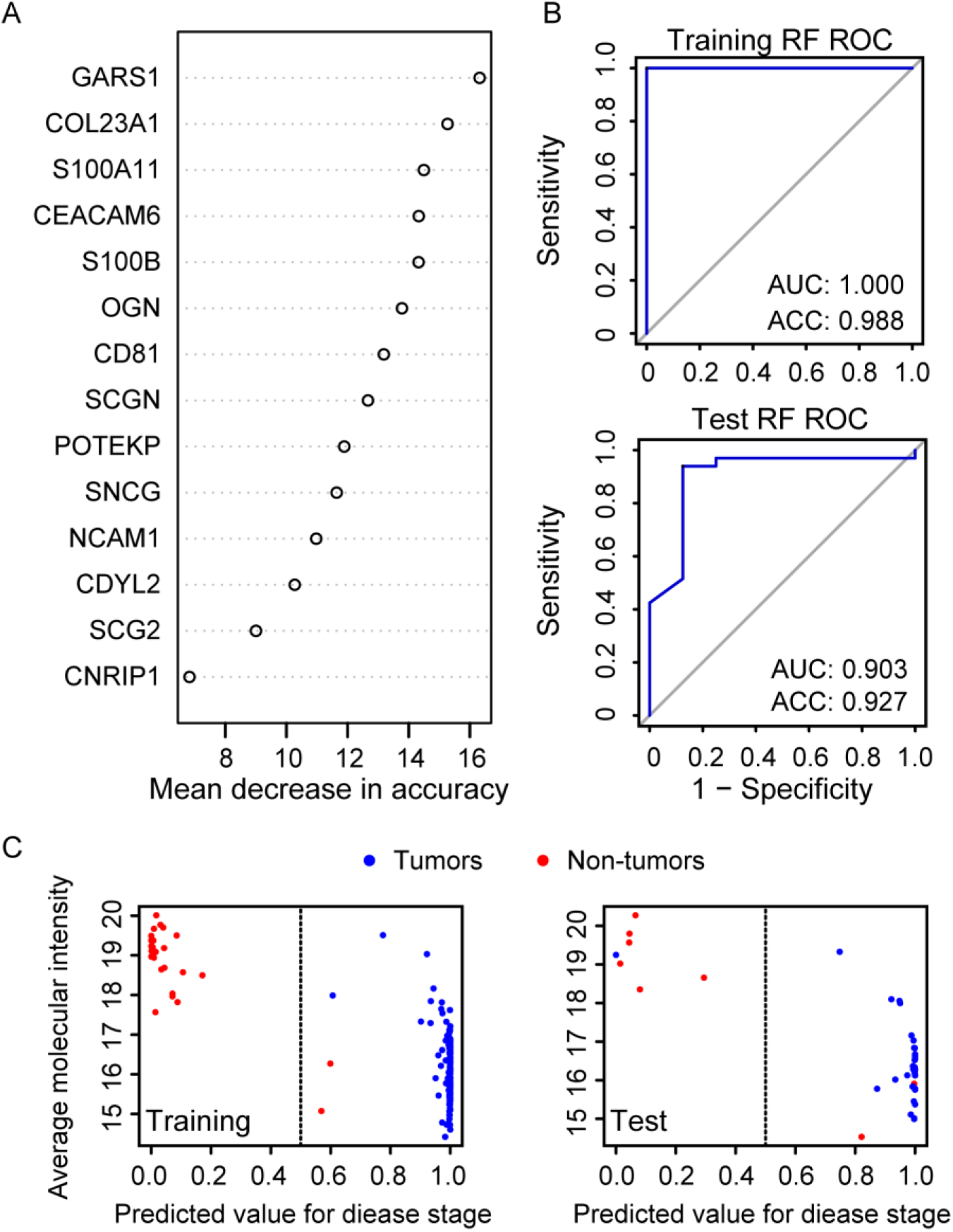
Machine learning to identify potential CRC biomarkers. (A) Prioritization of 14 important variables. (B) ROC plots for the training set (up) and the test set (down). (C) Performance of the model in the training set and test set.

### Analysis of protein isoforms and semi-tryptic peptides

We next checked whether this resource could be used to analyze specific protein isoform. Among the dysregulated proteins from IF, we identified SPTBN1 (SPTBN1-long) and one of its isoforms (SPTBN1-short)^43^. As reported in literature, SPTBN1 is significantly dysregulated and plays an essential role in liver cancer^44^, colorectal cancer, and breast cancer, among others^45, 46^. To assess the accuracy of the identification, we showed the sequence of SPTBN1-long and SPTBN1-short identified in the library, in addition to the common parts of the two sequences, our library had also identified the peptide (TSSISGPLSPAYTGQVPYNYNQLEGR) specific in SPTBN1-short (Figure 6A). The Skyline software (Skyline-daily version) was used to show the peak spectrum of this peptide and a common peptide form these two proteins within the DIA raw file (Figure 6B-C).

**Figure 6.**
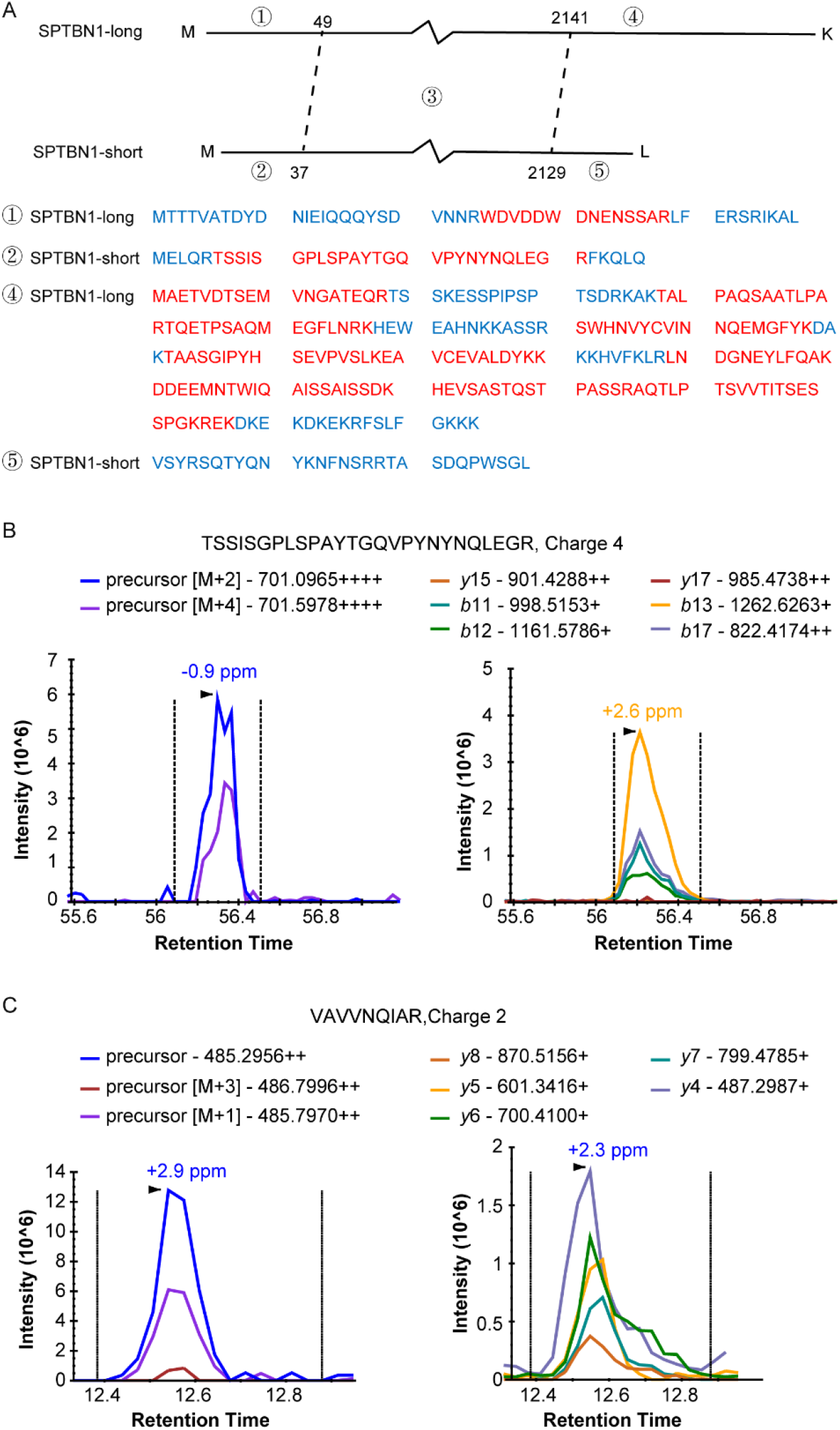
SPTBN1 protein identification in our DIA search results. (A) Sequences of SPTBN1 and its isoform. Blue: sequences that were not identified; red: identified sequences. (B) The peak spectrum of peptide SSISGPLSPAYTGQVPYNYNQLEGR in our DIA raw file (obtained using Skyline).

Regarding those were only characterized through semi-specific peptides in our semi-specific libraries (IS and RS), including VWF, LMO7, ALDH2, NPEPL1, NUAK1, and TPT1, many of them have important biologic implications. ADAM22 is a new therapeutic option for treating metastatic brain disease and may be appropriate for treatment of breast cancer^47, 48^. By analyzing mRNA expression profiles, Xin et al. found that *ASPM* is highly expressed in GBM, and patients with high *ASPM* expression have poor prognoses^49^. LRP6 inhibits cell proliferation and delays tumor growth in vivo, especially in colon, liver, breast, and pancreatic cancers^50, 51^. CHD9 was reported as a potential biomarker for clear cell renal cell carcinoma^52^. In addition, FAIM2 promotes non-small cell lung cancer growth and bone metastasis formation by regulating the epithelial-mesenchymal transformation process and the Wnt/β-catenin signaling pathway^53^. In our analysis, all these proteins showed significant differences between tumor and non-tumor samples, indicating that DPHL v2 can assist with the discovery of new potential protein biomarkers.

## Conclusion

We present DPHL v2: four comprehensive spectral libraries (RF, RS, IF, and IS) derived from 1608 DDA MS raw files, including 24 sample types. By identifying over 440,000 peptides and more than 14,000 proteins, DPHL v2 can confidently detect and quantify more than 66.1% of the reviewed human proteins annotated by UniProtKB/Swiss-Prot. Our results suggest that DPHL v2 could support protein biomarker identification, especially for protein isoforms and semi-tryptic peptides. DPHL v2 outperforms previous DIA libraries in the following aspects. Firstly, five additional tissue types (oral cavity, thymus, esophagus, eyelid, and ovary) and one blood plasma sample from T-ALL were included. Secondly, protein isoforms and semi-trypsin digestion were used for library searching. In addition, these libraries are compatible with various commonly used DIA tools, with or without format transformation, such as OpenSWATH^54^, DIA-NN^55^, Skyline^56^, and Spectronaut^57^.

## Materials and Methods

All chemicals used in this study were purchased from Sigma. All MS-grade reagents were acquired from Thermo Fisher Scientific (Waltham, MA).

### Clinical samples

Formalin-fixed paraffin-embedded, fresh or fresh frozen tissue biopsies from GBM, healthy human brain, eyelid tumor, thyroid disease, sarcoma, OSCC, thymus, LUAD, TNBC, HCC, gastric cancer, diffuse large B-cell lymphoma, pancreatic ductal adenocarcinoma, bladder cancer, PCa, and OV were collected in this study. Human plasma samples, including acute lymphoblastic leukemia (ALL), AML, T-ALL, normal plasma exosome, and blood disease, were also analyzed, as well as K562 cells. Six of these tissues were new additions compared to the DPHL v1. Eyelid samples were obtained from the Second Affiliated Hospital of Zhejiang University School of Medicine, China. The ovary cohort was obtained from The Cancer Hospital of the University of Chinese Academy of Sciences. The OSCC, esophagus, T-ALL, and thymus cancer samples were collected at Amsterdam UMC/VU Medical Center, Amsterdam, and Erasmus University Medical Center. Sample details are provided in Table S1.

To compare our libraries with the DPHL v1 and library-free method, we used the DIA data of a CRC cohort generated by Ge et al.^27^, which consists of 201 cancer samples, 40 para-cancer tissues, and 45 biological and technical replicates from 40 CRC patients and four healthy controls. The detailed sample information is given in Table S2.

### MS Data acquisition

Among the newly added 586 DDA raw data files, 108 were derived from Dutch cohorts generated at the Jimenez lab and 404 from Chinese cohorts generated at the Guo lab. The pipeline for generating these DDA files coincided with that used for the DPHL v1. The DDA raw files were centroided and converted into mzXML using ProteoWizard^58^ (version 3.0.11579). Carbamidomethylation was set as fixed modification at cysteine residues; oxidation was set as variable modification at methionine residues.

### DIA data analysis

The DIA raw files were submitted to DIA-NN (1.7.15), a tool for DIA or SWATH proteomics data analysis^55^. Our four libraries were used as a reference, and no other fasta sequences were added. The library inference was set to “off”. All other parameters were kept to their default values. The tools we used for the DIA data analysis, as described above, are publicly available^55^.

### Machine learning

The random forest analysis was performed with the R package “randomForest” (version 4.6.14). 1426 proteins were firstly selected as input features to build 1000 trees with 5-fold cross validation and repeated 10 times to optimize the model. The Mean Decrease Accuracy was set 4 to 6, with step size of 0.5. The final performance was evaluated by mean accuracy (ACC) and mean area under curve (AUC) in a receiver operating characteristic curve across 5-folds.

## Ethical statement

Ethics approvals for this study were obtained from the Ethics Committee or Institutional Review Board of each participating institution.

## Acknowledgments

This work is supported by grants from National Key R&D Program of China (2021YFA1301603; 2021YFA1301602; 2020YFE0202200).

## Author contributions

T.G. conceived the project. Z.X. and T.Z. built all the libraries. Z.X. processed and analyzed data. T.G., Z.X., T.Z., J.A., and F.Z. wrote the manuscript. Y.L. collected the brain samples. J.Y. provided the eyelid tumor samples. T.L. offered the lung cancer samples. J.Z. and C.L. collected the liver cancer samples. Y.H. offered the prostate cancer samples. Q.W. provided the cervix cancer samples. J.Z. and Z.Z. collected the ovarian cancer samples. Others prepared peptides for the study. T.G. supervised the work. All authors reviewed and approved the manuscript.

## Data Availability

All newly added raw DDA MS data, spectral libraries, and protein results are publicly available at iProX^59^ (PXD015314) and ProteomeXchange (PXD015314). All the R scripts were uploaded to GitHub (https://github.com/zhutiansheng/DPHLv2).

