## supporting information for "DPHL v2: An updated and comprehensive DIA pan-human assay library for quantifying more than 14,000 proteins"

Province, China
[11] Department of Urology, The Second Hospital of Dalian Medical University, No.467 Zhongshan Road, Dalian, Liaoning Province, China
[12] Department of Clinical Epidemiology, Shengjing Hospital of China Medical University, Shenyang 110000, Liaoning Province, China
[13] The Cancer Hospital of the University of Chinese Academy of Sciences (Zhejiang Cancer Hospital), Hangzhou, Zhejiang, China
[14] Institute of Basic Medicine and Cancer (IBMC), Chinese Academy of Sciences, Hangzhou, Zhejiang, China
[15] College of Mathematics and Computer Science, Zhejiang A & F University, Hangzhou, China.

**Figure S1. Correlation of paired peptides with iRT correction values before and after QC.**

**Figure S2. Characteristics of the RF library.**

**Figure S3. Characteristics of the RS library.**

**Figure S4. Characteristics of the IF library.**

**Figure S5. Characteristics of the IS library.**

**Figure S6. Bar plots displaying the number of identified proteins using our four libraries for each sample type.**

**Figure S7. Bar plots displaying the number of identified unique proteins for each sample type.**

**Figure S8. SubLib DIA analysis of the tissue samples from CRC and benign samples.**

**Table S1. Summary information of the tissue samples used for the DPHLv2 project.**

**Figure S1.**

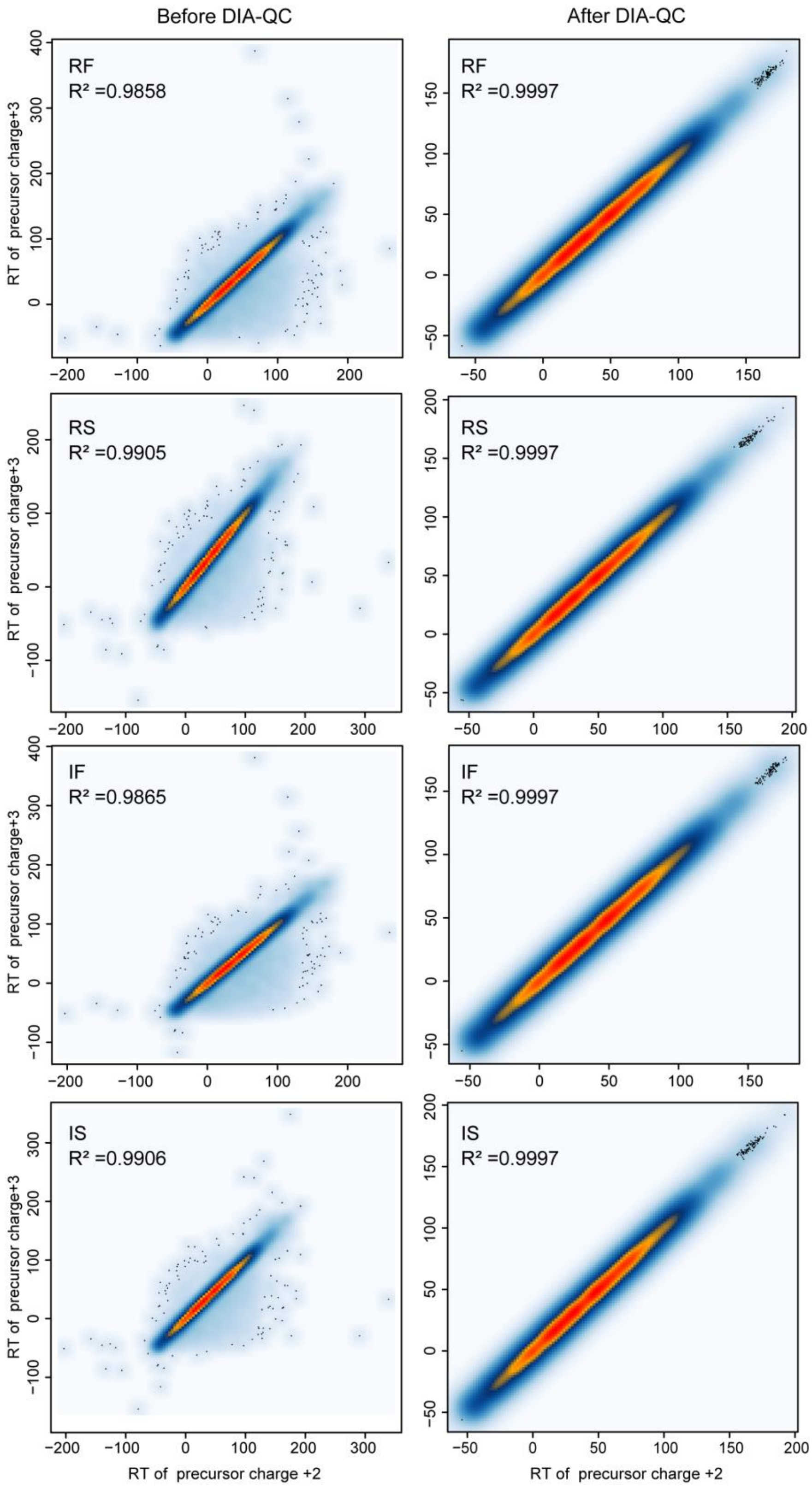


**Figure S1. Correlation of paired peptides with iRT correction values before and after QC.**

**Figure S2.**

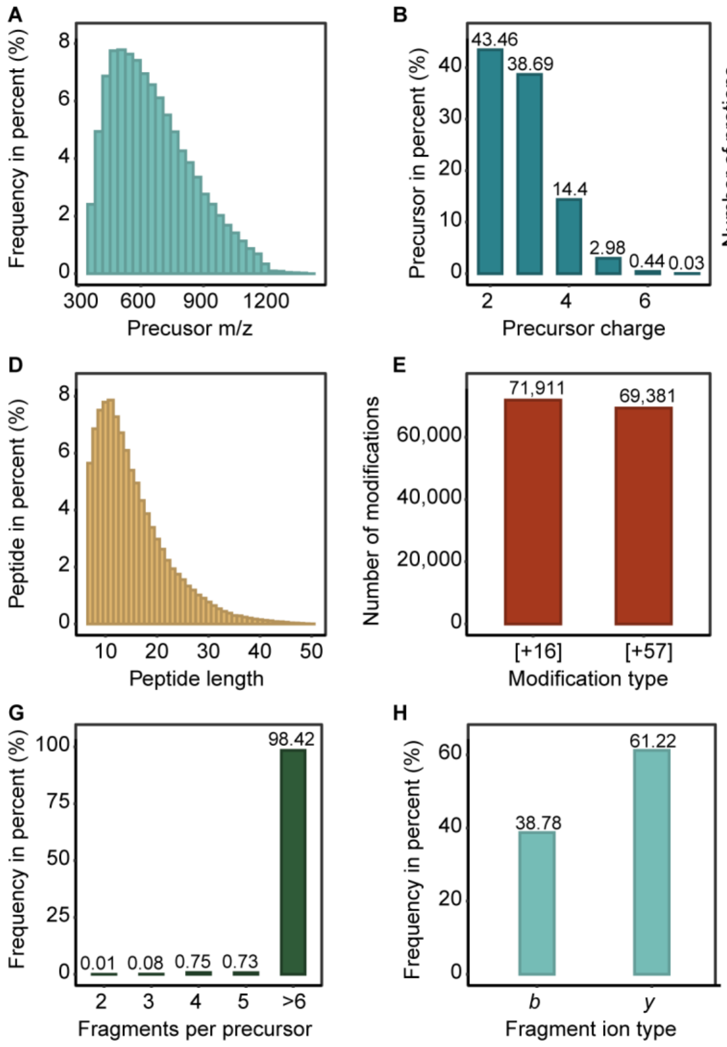


**Figure S2. Characteristics of the RF library.** (A) Distribution of precursors' m/z. (B) Counts of different precursor charge states. (C) Number of proteotypic peptides for each protein. (D) Distribution of peptide lengths. (E) Number of peptides with either of two modifications (+16: oxidation in methionine; +57: carbamidomethylation in cysteine). (F) The proportion of different charges of fragment ions. (G) Proportion of fragment ions per precursor ion. (H) Percentage of b and y ions.

**Figure S3.**

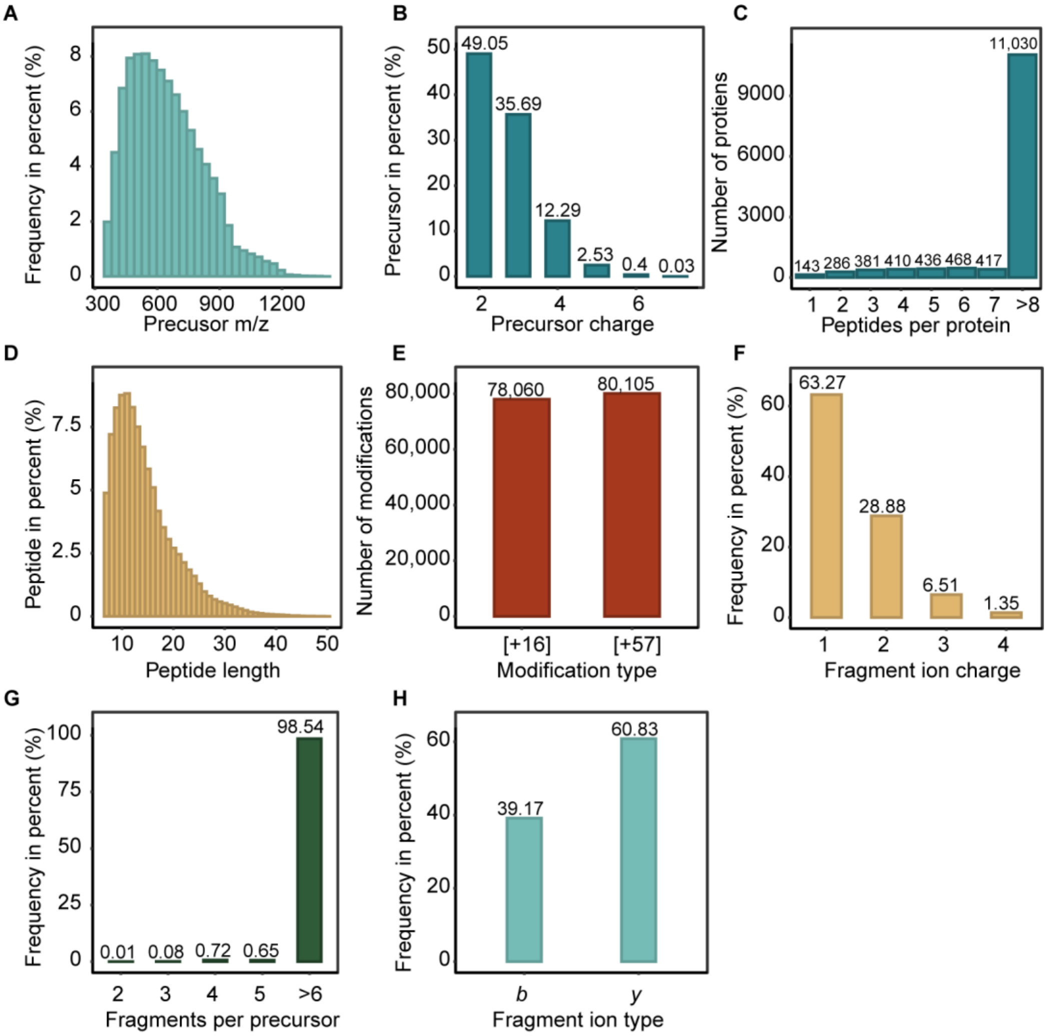


**Figure S3. Characteristics of the RS library.** (A) Distribution of precursors' m/z. (B) Counts of different precursor charge states. (C) Number of proteotypic peptides for each protein. (D) Distribution of peptide lengths. (E) Number of peptides with either of two modifications (+16: oxidation in methionine; +57: carbamidomethylation in cysteine). (F) The proportion of different charges of fragment ions. (G) Proportion of fragment ions per precursor ion. (H) Percentage of b and y ions.

**Figure S4.**

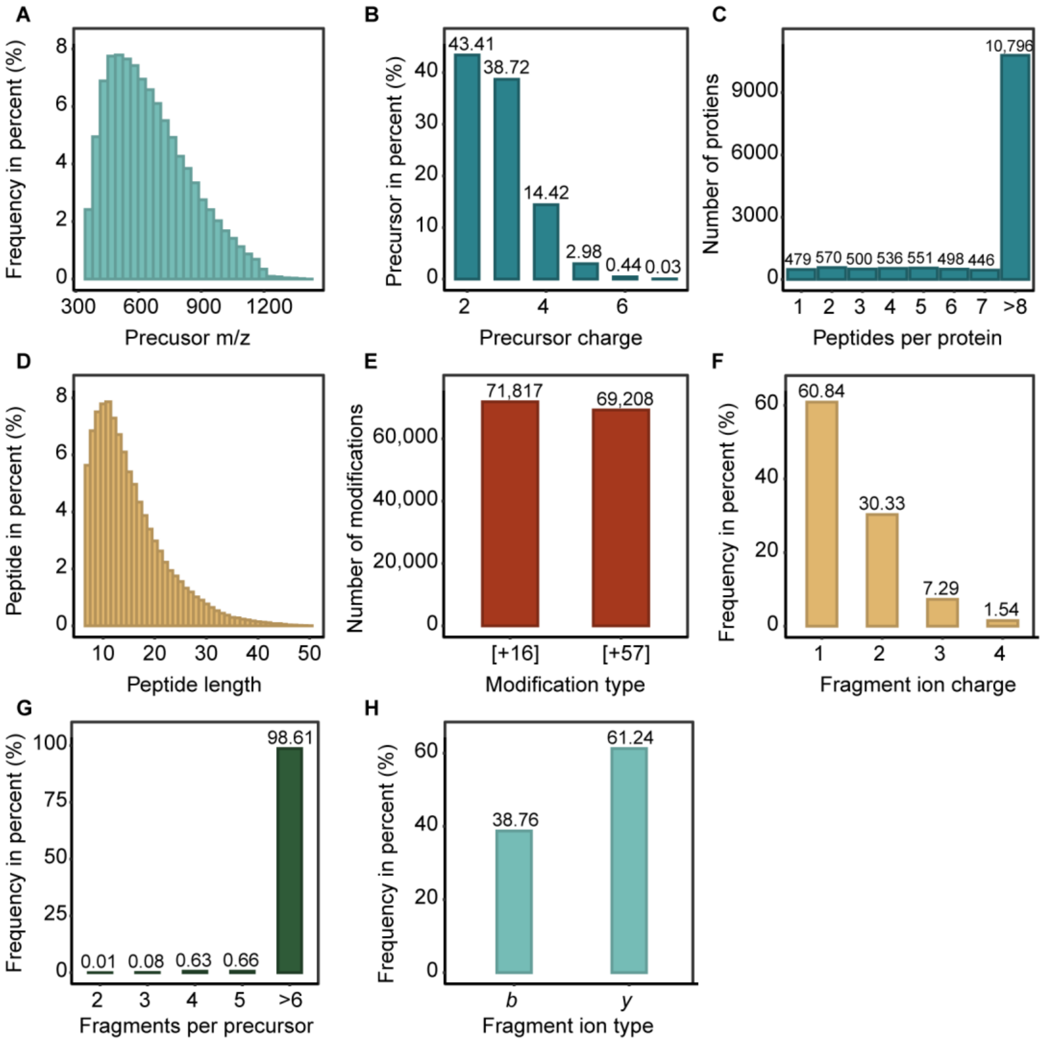


**Figure S4. Characteristics of the IF library.** (A) Distribution of precursors' m/z. (B) Counts of different precursor charge states. (C) Number of proteotypic peptides for each protein. (D) Distribution of peptide lengths. (E) Number of peptides with either of two modifications (+16: oxidation in methionine; +57: carbamidomethylation in cysteine). (F) The proportion of different charges of fragment ions. (G) Proportion of fragment ions per precursor ion. (H) Percentage of b and y ions.

**Figure S5.**

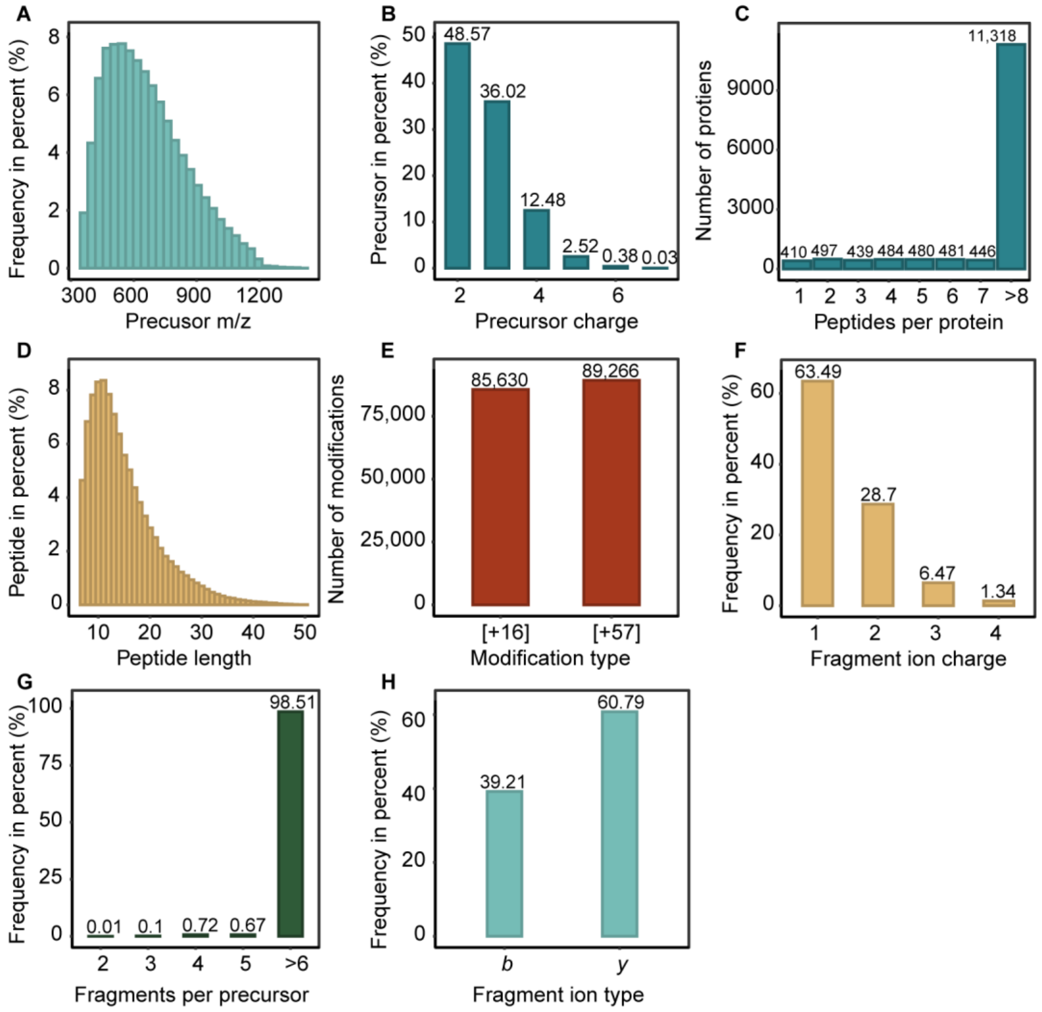


**Figure S5. Characteristics of the IS library.** (A) Distribution of precursors' m/z. (B) Counts of different precursor charge states. (C) Number of proteotypic peptides for each protein. (D) Distribution of peptide lengths. (E) Number of peptides with either of two modifications (+16: oxidation in methionine; +57: carbamidomethylation in cysteine). (F) The proportion of different charges of fragment ions. (G) Proportion of fragment ions per precursor ion. (H) Percentage of b and y ions.

**Figure S6.**

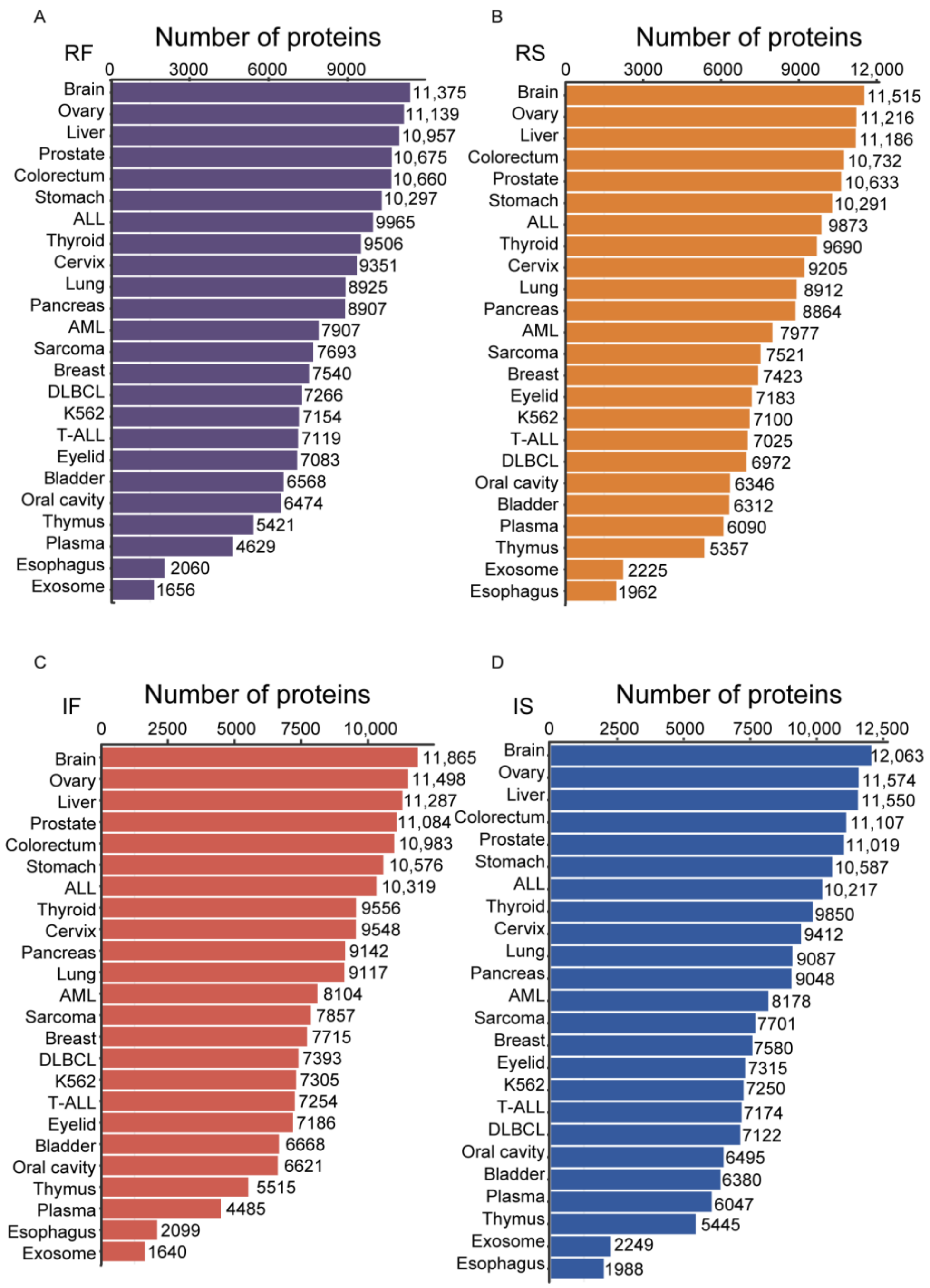


**Figure S6. Bar plots displaying the number of identified proteins using our four libraries for each sample type.** RF, reviewed fasta sequence & full-specific digestion mode; RS, reviewed fasta sequence & semi-specific digestion mode; IF, isoform fasta sequence & full-specific digestion mode; IS, isoform fasta sequence & semi-specific digestion mode.

**Figure S7.**

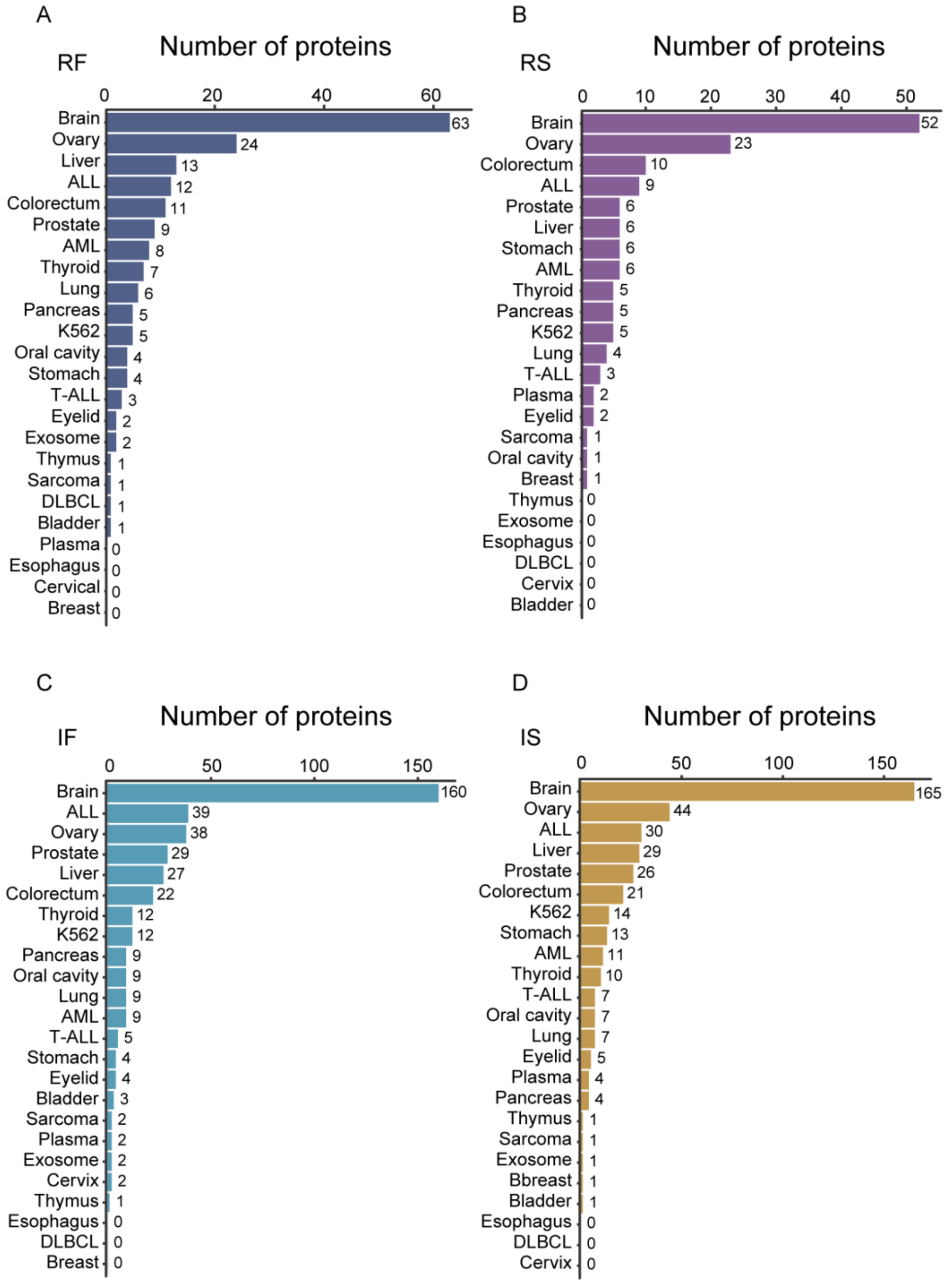


**Figure S7. Bar plots displaying the number of identified unique proteins for each sample type.** RF, reviewed fasta sequence & full-specific digestion mode; RS, reviewed fasta sequence & semi-specific digestion mode; IF, isoform fasta sequence & full-specific digestion mode; IS, isoform fasta sequence & semi-specific digestion mode.

**Figure S8.**

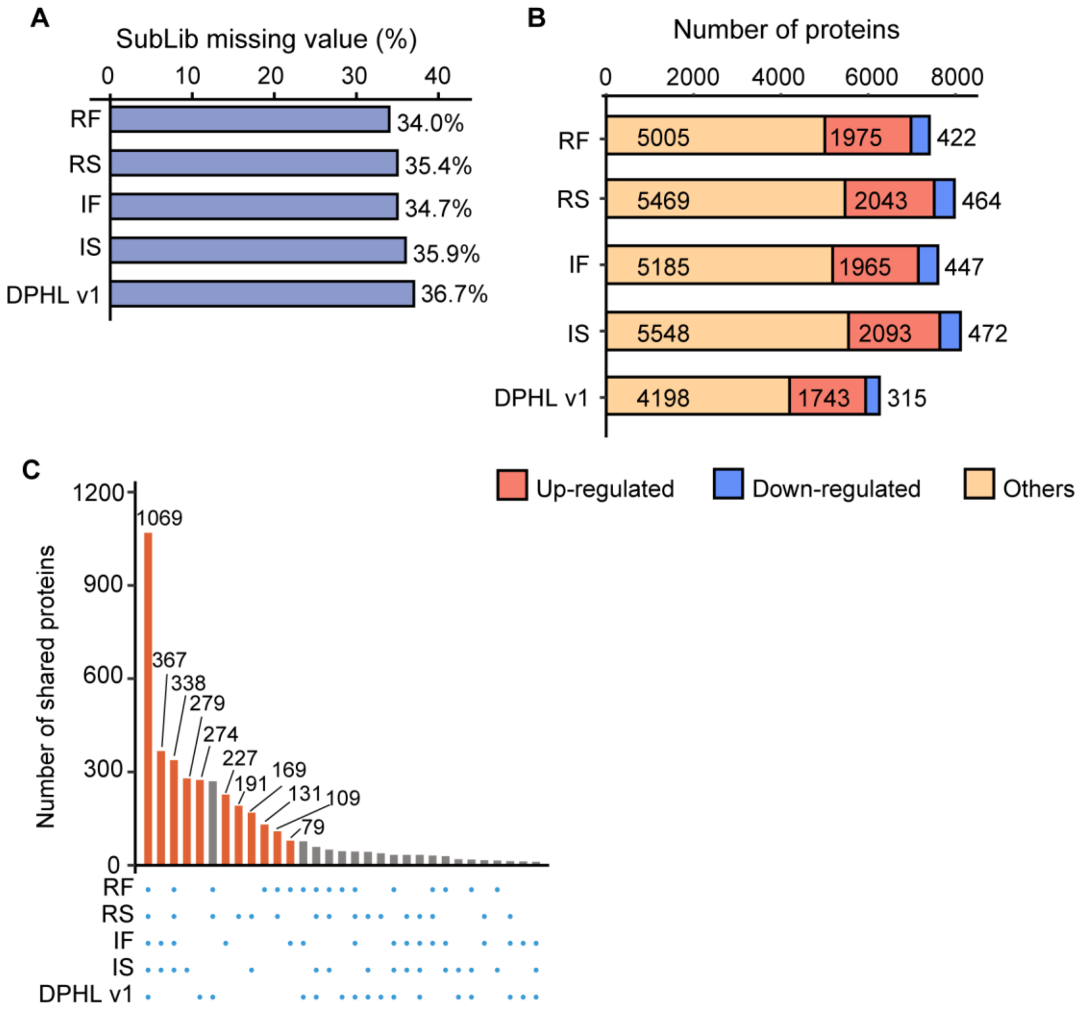


**Figure S8. SubLib DIA analysis of the tissue samples from CRC and benign samples.** (A) Missing values derived from the five libraries. (B) Number of differentially expressed proteins between CRC and benign samples using the five libraries. Proteins with adjust p value < 0.01 and log2 (fold-change) > 1 were selected as significantly differentially expressed. FC, fold change. (C) Protein identification overlaps across the five libraries.
